# PTBP1 mediates Sertoli cell actin cytoskeleton organization by regulating alternative splicing of actin regulators

**DOI:** 10.1101/2024.06.12.598725

**Authors:** Yuexi Wang, Ullas Valiya Chembazhi, Danielle Yee, Sijie Chen, Jie Ji, Yujie Wang, Ka Lam Nguyen, PoChing Lin, Antonia Ratti, Rex Hess, Huanyu Qiao, CheMyong Ko, Jing Yang, Auinash Kalsotra, Wenyan Mei

## Abstract

Spermatogenesis is a biological process within the testis that produces haploid spermatozoa for the continuity of species. Sertoli cells are somatic cells in the seminiferous epithelium that orchestrate spermatogenesis. Cyclic reorganization of Sertoli cell actin cytoskeleton is vital for spermatogenesis, but the underlying mechanism remains largely unclear. Here, we report that RNA-binding protein PTBP1 controls Sertoli cell actin cytoskeleton reorganization by programming alternative splicing of actin cytoskeleton regulators. This splicing control enables ectoplasmic specializations, the actin-based adhesion junctions, to maintain the blood-testis barrier and support spermatid transport and transformation. Particularly, we show that PTBP1 promotes actin bundle formation by repressing the inclusion of exon 14 of *Tnik*, a kinase present at the ectoplasmic specialization. Our results thus reveal a novel mechanism wherein Sertoli cell actin cytoskeleton dynamics is controlled post-transcriptionally by utilizing functionally distinct isoforms of actin regulatory proteins, and PTBP1 is a critical regulatory factor in generating such isoforms.

## INTRODUCTION

Spermatogenesis is a complex physiological process that produces haploid spermatozoa for the continuity of species. Spermatogenesis occurs within the seminiferous tubules of the testis. During spermatogenesis, spermatogonial stem cells undergo mitotic division to self-renew and generate preleptotene spermatocytes (1). Subsequently, via meiosis, spermatocytes progress into haploid spermatids. This is followed by spermiogenesis, wherein round spermatids undergo a series of transformation steps and become elongated spermatids. This process encompasses a sequence of events, such as the condensation of genetic material, flattening of the acrosome, disposal of the residual body, and the formation of the flagellum (1). Lastly, elongated spermatids are released from the seminiferous epithelium and translocate to the epididymis for final maturation, a phenomenon referred to as spermiation. Impaired spermatogenesis and spermiation reduce the sperm number and quality, leading to male infertility.

Sertoli cells, the only somatic cell lineage within the seminiferous epithelium, have been reported to orchestrate almost every aspect of spermatogenesis (2). Sertoli cells not only provide biomolecules such as niche signals and micronutrients to germ cells but also structurally support their translocation, positioning, transformation, and release (2). The supporting role of Sertoli cells relies on the remarkable dynamics of Sertoli cell cytoskeleton. Actin microfilaments, microtubules, and intermediate filaments are three major elements of Sertoli cell cytoskeletons. They concentrate at specific regions of Sertoli cells and function together to support germ cell development (3). Among them, filamentous actin (F-actin) is a critical component of the ectoplasmic specialization (ES), one type of specialized adhesion junctions formed by Sertoli cells for intercellular attachment between Sertoli-Sertoli cells and Sertoli-germ cells(3).

ESs are found at both the apical and basal regions of Sertoli cells. At the basal region, ESs of two neighboring Sertoli cells intimately associate with tight junctions, gap junctions, and desmosomes to form the blood-testis barrier (BTB) (3,4). The BTB segregates seminiferous tubules into the basal and adluminal compartments. This allows the differentiation of spermatogonia up to the preleptotene stage to occur at the basal compartment, while meiosis, spermiogenesis, and spermiation take place at the adluminal compartment (4). During the seminiferous epithelium cycle, the BTB undergoes periodical disassembly and reconstruction, allowing the preleptotene spermatocytes to translocate from the basal to the adluminal compartment for meiosis (5,6). As such, immunogenic meiotic and post-meiotic germ cells are sequestered in the immune-privileged adluminal compartment and protected from autoimmune attacks. At the apical region, ESs form at the Sertoli cell regions around the heads of differentiating spermatids to anchor spermatids and maintain their polarity. This physical arrangement of spermatids facilitates the cellular interactions between spermatids and Sertoli cells during spermiogenesis and maximizes the usage of the seminiferous epithelium space, allowing millions of spermatozoa to be produced in a day. Similar to the basal ES, the apical ES undergoes cyclic breakdown and reconstruction to enable spermatid transport across the adluminal compartment, and the elongated spermatids line up near the luminal edge for spermiation (6).

Each ES consists of a layer of hexagonally packed actin filament bundles that are sandwiched between the attached cistern of the endoplasmic reticulum and the nearby Sertoli cell plasma membrane(3). The actin bundles have been reported to reinforce the intercellular adhesion domains of the ES (7,8). As such, actin bundles in the ES must form and disassemble following the progression of a seminiferous epithelial cycle to enable rapid ES turnover (9–11). A number of actin regulatory proteins, including some required for actin assembly, have been found present at the ES (12–27). Yet, the molecular mechanism by which actin cytoskeleton reorganization is timely controlled in Sertoli cells remains poorly understood.

Alternative splicing (AS) is an important post-transcriptional regulatory mechanism that increases the transcriptome and proteome complexity. By selectively joining alternative exons from a single gene, AS produces multiple mRNA splice variants from the same gene and consequently affects the mRNA stability, localization, or translation, rapidly altering the protein landscape of a cell. Notably, Sertoli cells exhibit a very complex AS pattern that generates more than 45% novel protein isoforms of known genes (28). How AS in Sertoli cells contributes to their function in Spermatogenesis is unclear.

The polypyrimidine tract binding protein 1 (PTBP1, also known as hnRNP I) is an RNA-binding protein known for generating cell type-specific AS (29). In the seminiferous tubules, PTBP1 is predominantly expressed in Sertoli cells and mitotic spermatogonia cells (30–33). Deletion of PTBP1 in mouse germ cells suppresses spermatogonia cell proliferation, resulting in male infertility (32). Knocking down PTBP1 in the TM4 Sertoli cell line *in vitro* reduces the expression of tight junction proteins and increases the BTB permeability (33). However, it was unclear how PTBP1 controls AS in Sertoli cells to support spermatogenesis *in vivo*. Here, we report that deletion of PTBP1 in mouse Sertoli cells disrupts the BTB function, spermatid transport, adhesion, and differentiation, resulting in male infertility. Actin cytoskeleton organization at both the basal and apical ES is impaired. Mechanistically, we show that PTBP1 regulates AS of a number of actin cytoskeleton regulators in Sertoli cells. Of particular, PTBP1 represses the inclusion of exon 14 of Traf2 and Nck-interacting kinase (TNIK) to promote actin bundle formation in Sertoli cells. Our findings, thus, uncover a novel mechanism wherein PTBP1 regulates the AS of actin cytoskeleton regulators in Sertoli cells to orchestrate spermatogenesis.

## MATERIALS AND METHODS

### Animals

#### Generation of *Ptbp1*^flox/flox^; *Amh-Cre* mice

The *Ptbp1*^flox/flox^ allele (*Ptbp1*^fl/fl^) was previously reported (34). The floxed *Ptbp1* mice were crossed with the *Amh-cre* mice (JAX stock #007915) (35) to generate the *Ptbp1*^fl/fl^; *Amh-Cre*^+/-^ mice. All mice used in this report were from the breeding cross of the *Ptbp1*^fl/fl^; *Amh-Cre* female mice with the *Ptbp1*^fl/fl^ male mice. The *Ptbp1*^fl/fl^; *Amh-Cre* mice were referred to as the Sertoli cell-specific *Ptbp1* knockout mice (*Ptbp1*^ΔSC^), and their littermates *Ptbp1*^fl/fl^ mice were referred to as the control mice. Primer sequences for genotyping mice are shown in Supplementary Table 1.

### Ethics statement of animal use

All mice used in these experiments were housed in the College of Veterinary Medicine at the University of Illinois Urbana-Champaign (UIUC) and cared for according to the institutional “Guide for the Care and Use of Laboratory Animals”. All procedures involving mouse care, euthanasia, and tissue collection were approved by the UIUC Animal Care and Use Committee (IACUC approved protocol #23096).

### Testis weight measurement

The weights of the left and right testes of each mouse were measured and averaged (mg).

### Histology and immunostaining

Dissected testes were fixed in Bouin’s fixative or 4% paraformaldehyde (PFA), paraffin-embedded and sectioned according to the standard protocols. Testis sections (5 μm) were processed for histological staining or immunostaining. The Periodic acid-Schiff (PAS) and hematoxylin staining was performed according to standard protocols. For immunofluorescence staining, deparaffinized sections were heated in sodium citrate buffer (10 mM sodium citrate, 0.05% Tween-20, pH 6.0) for 20 minutes (min) for antigen retrieval. After 1 hour (h) of blocking (5% serum, 0.1% TritonX-100) at room temperature, sections were incubated with primary antibodies at 4 ^°^C overnight. After washing with PBST, sections were incubated with fluorescence-conjugated secondary antibodies for 2 h at room temperature. 4′,6-diamidino-2-phenylindole (DAPI) was used for staining nuclei. For phalloidin staining, testes were fixed in 4% PFA, embedded in OCT, and cryosectioned. Frozen testis sections were incubated with AlexaFluor 488-conjugated phalloidin for 15 min at room temperature. For immunohistochemistry, R.T.U vectastain kit (Vector Laboratories) and DAB substrate were used, and sections were lightly counterstained with hematoxylin to label cell nuclei. Images were taken from a Leica or Keyence compound microscope or a Nikon A1R confocal microscope and processed using Fiji.

#### Antibodies

Secondary antibodies used were AlexaFluor 488- or 594-conjugated goat anti-rabbit, mouse, or rat and AlexaFluor 488- or 594-conjugated donkey anti-goat or rabbit. Primary antibodies used were rabbit anti-GATA4 (Cell Signaling Technology, Cat# 36966), rat anti-GATA4 (Thermo Fisher Scientific, Cat# 14-9980-80), mouse anti-PLZF (Santa Cruz Biotechnology, Cat# sc-28319), rabbit anti-PTBP1 (Cell Signaling Technology, Cat# 72669), goat anti-PTBP1 (Novus Biologicals, Cat# NB-100-1310), rat anti-ZO1 (DSHB, #R26.4C), rabbit anti-CLAUDIN 11 (Thermo Fisher Scientific, Cat#36-4500), mouse anti-ESPIN (ESPN) (BD Biosciences, #611656), rabbit anti-Androgen Receptor (Lab Vision, Cat# RB-9030-P0), mouse anti-TUBB3 (Thermo Fisher Scientific, Cat# MA1-19187), rabbit anti-TNIK (Invitrogen, # PIPA120639), mouse anti-phospho-HISTONE H2A.X (Ser139) (γH2AX) (Sigma-Aldrich, Cat# ZMS05636), rabbit anti-SCP3 (Abcam, Cat# ab15093), rabbit anti-Cleaved CASPASE 3 (Cell Signaling Technology, Cat# 9661), rat anti-GCNA1 (DSHB, Cat# 10D9G11). Alexa Fluor 488-conjugated Phalloidin (Cell Signaling Technology, #8878) was used to stain F-actin.

### Fertility test

Sexually mature *Ptbp1*^fl/fl^ or *Ptbp1*^ΔSC^ males were bred with fertile *Ptbp1*^fl/fl^ females for over 6 months. The number of litters produced by each male was recorded and compared between *Ptbp1*^fl/fl^ and *Ptbp1*^ΔSC^ males.

### The BTB integrity assay

To analyze the integrity of the BTB, we used an assay based on the ability of an intact BTB to exclude the diffusion of a small molecule sulfo-NHSLC-biotin from the basal to the apical compartment of the seminiferous epithelium as previously reported (36). Briefly, 3 *Ptbp1*^ΔSC^ mice and 3 control *Ptbp1*^fl/fl^ mice at 3 months of age were anesthetized by isoflurane. Their testes were exposed by making small incisions of approximately 0.5 cm in the middle of the scrotum. 100 μl of EZ-Link™ sulfo-NHS-LC-biotin (at 10 mg/ml in PBS containing 1 mM CaCl2) stock was loaded gently under the tunica albuginea using a 28-gauge insulin syringe. After the sulfo-NHSLC-biotin administration, mice were allowed to rest for 30 min under anesthesia and then euthanized. Testes were collected, embedded in OCT, and cryosectioned to get 7 μm-thick frozen sections. Testis frozen cross sections were then fixed at room temperature in 4% PFA for 10 min, washed with PBS, and stained with Alexa Fluor 555-streptavidin for 30 min at room temperature. After staining, sections were washed with PBS and stained with DAPI to label cell nuclei. Images were taken on a Nikon A1R confocal microscope.

### Transmission electron microscopy (TEM)

3 *Ptbp1*^ΔSC^ and 2 *Ptbp1*^fl/fl^ male mice at P100 were perfused with PBS and a fixation buffer (0.1 M Na-Cacodylate, 2% PFA, and 2.5% glutaraldehyde (pH 7.4)). After perfusion, testes were dissected and fixed in the aforementioned fixation buffer at 4°C overnight. The testes were then washed with 0.1 M Na-Cacodylate buffer and post-fixed in 0.1 M Na-Cacodylate buffer containing 1% aqueous osmium tetroxide (pH 7.4). After rinsed in 0.1 M Na-Cacodylate buffer, testes were stained in 2% aqueous uranyl acetate at 4 °C overnight, dehydrated using a gradient of ethanol, sequentially infiltrated with 1:1 and 1:2 of ethanol: propylene oxide, 1:1 and 1:2 of propylene oxide: Polybed 812 mixture. Finally, testes were embedded in a 100% Polybed 812 mixture containing 1.5% DMP-30 at 60 °C for 24 h. Then, 100 nm-thick sections were prepared and placed on PELCO 200 Mesh grids. Images were acquired using the Thermo Fisher FEI Tecnai G2 F20 S-TWIN STEM with an AMT 4k x 4k CMOS camera under 160kV.

### The PCR-based splicing assay

Primers were designed to target the constitutive exons that flank the alternative exons of interest. Primer sequences are shown in Supplementary Table 2. The PCR amplicons were resolved on the 3% agarose gel containing ethidium bromide and imaged using the Carestream MI SE system. Quantification of gel images was done by Fiji (NIH). Percentage spliced in values (PSI) were determined as the exon inclusion band intensity/(the exon inclusion band intensity + the exon exclusion band intensity) x 100.

### RNA-seq analysis

RNAs were extracted from whole testes of 3 *Ptbp1*^fl/fl^ and 3 *Ptbp1*^ΔSC^ mice at postnatal day (P) 35 using the PureLink RNA Mini Kit (Ambion, Cat. 12183025). RNA-seq libraries were constructed using Illumina’s TruSeq Stranded mRNA sample prep kit, pooled and sequenced on one SP lane for 151 cycles from both ends of the fragments on a NovaSeq 6000 at the UIUC Biotechnology Center High-Throughput Sequencing Core. Each library generated over 160 million paired reads. Raw reads were subjected to read length and quality filtering using Trimmomatic V0.38 (37) and aligned to the mouse genome (mm10) using STAR (version 2.6.1d) (38). Cufflinks package (39) was used to assess differential gene expression events, among which significant events were identified using a stringent cutoff criterion: FDR(q-value) < 0.05, FPKM≥1, and −1.5 ≥ fold change≥1.5. rMATS v4.0.2(turbo)(40) was used to study differential splicing, and events with FDR < 0.1, junction read counts ≥ 10, and PSI ≥ 10% were deemed to be significant. Exon ontology analysis was performed on the set of alternatively spliced cassette exons identified using rMATS. Mouse (mm10) annotations were converted to human (hg19) annotations using UCSC liftover with a minimum base remap ratio set to 0.8. Exon ontology pipeline (41) was then used on the lifted exons to perform ontology analysis.

### Sertoli cell isolation and culture

Primary Sertoli cells were cultured following the published protocol with a few modifications (42). In brief, testes were isolated from P22 or P35 mice. An incision was made on the tunica albuginea to collect seminiferous tubules. Tubules were gently loosened and incubated in the digestion solution 1 containing 1 mg/mL Collagenase Type1 (Gibco) and 20 μg/mL DNase I (Roche) in DMEM/F12 and shaken at 37 °C for 30 min. Afterward, tubules were transferred to the digestion solution 2 containing 1 mg/mL Collagenase Type1, 20 μg/mL DNase I, and 2 mg/mL hyaluronidase (MP Biomedicals) in DMEM/F12 and shaken at 37 °C for 15 min. The fragmented tubules were spun down at 100 g for 1 min and incubated in the fresh digestion solution 2. After another 20 min of shaking, cells were centrifuged down, and the enzyme digestion was stopped by washing cells several times in Sertoli cell culture media (DMEM/F12 containing 10% fetal bovine serum, 1x Penicillin/Streptomycin, and 3 mM L-Glutamine). Finally, cells were resuspended in the Sertoli cell culture medium and seeded in 24-well plates. For immunocytochemistry, isolated Sertoli cells were seeded on poly-L lysine-coated coverslips (Corning) in 24-well plates and cultured at 37 ^°^C in a humidified incubator supplied with 5% CO_2_. After 48 h of culture, cells were treated with hypotonic solutions containing 20 mM 2-Amino-2-hydroxymethylpropane-1,3-diol (TRIS)-HCl at pH 7.5 for 2.5 min at room temperature to remove germ cells. Sertoli cell culture media was changed every 2 days.

### Splicing-inhibiting morpholino antisense oligonucleotides (ASO) transfection

Morpholino ASO complementary to the 3’ end of *Tnik* exon 14 and the adjacent 3’ splice site in the intron region was designed to block the inclusion of exon 14 and ordered from Gene Tools. The control ASO was a standard control oligo ordered from Gene Tools. Sequences of *Tnik* ASO and control ASO are shown in Supplementary Table 3. Sertoli cells were isolated and purified from P22 *Ptbp1*^fl/fl^ and *Ptbp1*^ΔSC^ testes as described above. When Sertoli cells reached 60% confluency, they were transfected with 3 μM *Tnik* ASO, control ASO, or sterile water (as the vehicle control) using 6 μM Endo-Porter as the manufacturer suggested. 48 h after the transfection, Sertoli cells were either fixed in 4% PFA for the immunocytochemistry staining or harvested for the PCR-based splicing assay.

### Statistics analysis

Differences between the *Ptbp1*^fl/fl^ and *Ptbp1*^ΔSC^ mice were assessed for significance using an unpaired Student’s 2-tailed *t*-test unless otherwise noted. Data involving three or more variables were analyzed by one-way ANOVA and Turkey’s *post hoc* test using GraphPad Prism.

## RESULTS

### Loss of PTBP1 in Sertoli cells results in male infertility

To determine the function of PTBP1 in Sertoli cells, conditional deletion of PTBP1 in Sertoli cells was accomplished by generating *Ptbp1*^fl/fl^; *Amh-Cre*^+/-^ (hereafter *Ptbp1*^ΔSC^) mice starting during embryogenesis (35). Testes of *Ptbp1*^ΔSC^ mice were significantly smaller at P25 compared to those of their sibling littermate control *Ptbp1*^fl/fl^ mice and this difference became more prominent in older mice (Figure 1A and 1B). To evaluate fertility, *Ptbp1*^ΔSC^ males were bred with wild-type fertile females. On average, *Ptbp1*^ΔSC^ males produced significantly fewer litters over 6 months, and 33% never gave any pups throughout their reproductive age (Figure 1C), suggesting that PTBP1 function in Sertoli cells is essential for male fertility.

**Figure 1.**
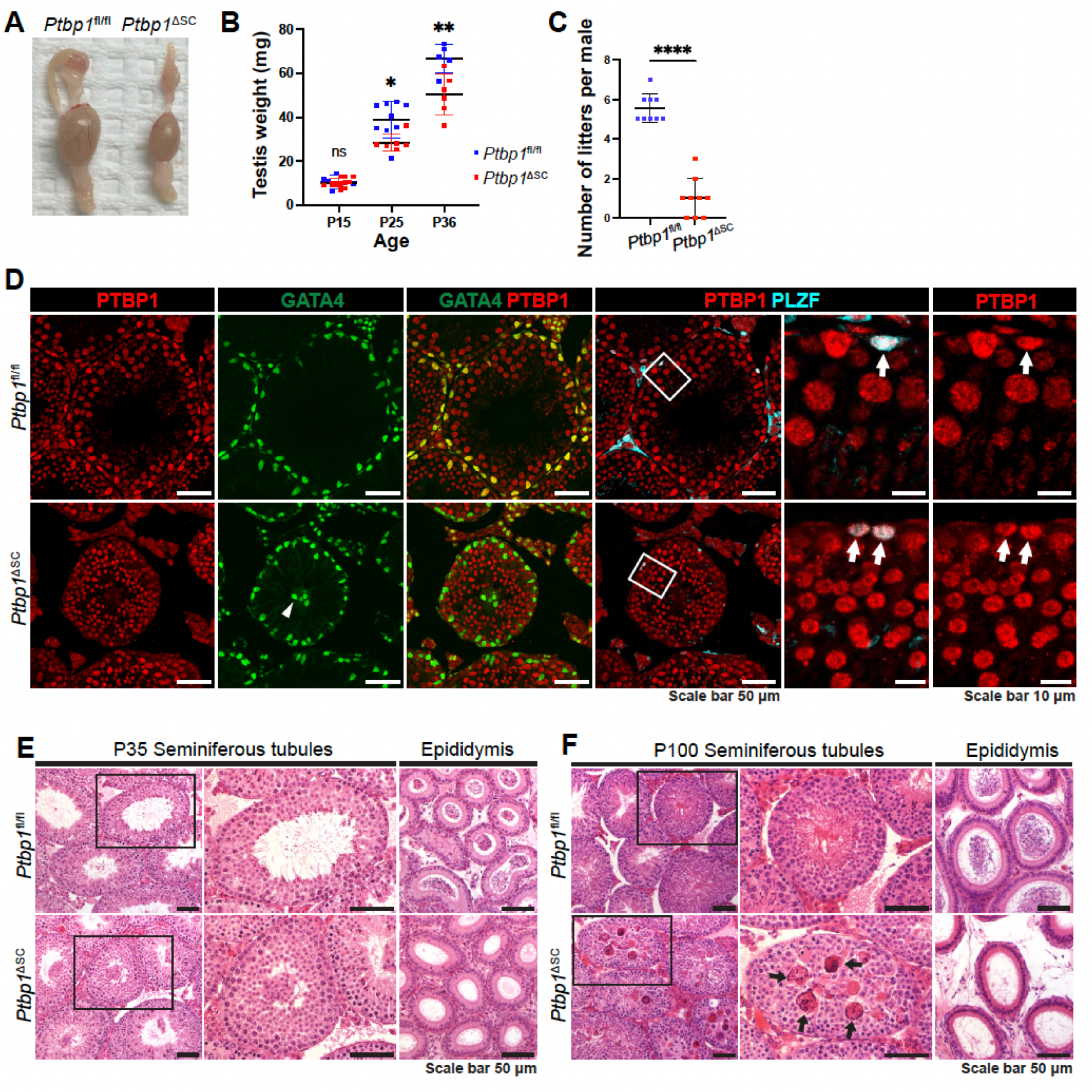
Sertoli cell-specific PTBP1 deletion disrupted spermatogenesis. (A-B) shows the size difference between the testes from *Ptbp1*^f/f^ and *Ptbp1*^ΔSC^ mice. (C) shows the number difference between the litters generated by *Ptbp1*^f/f^ and *Ptbp1*^ΔSC^ mice. Data in B and C are presented as mean ± SD. NS: not significant. ***P<0.001, **** p < 0.0001. Each symbol in B and C represents the value of each mouse. (D) Double immunofluorescence staining using antibodies against PTBP1, GATA4 (a marker for Sertoli cells), or PLZF (a marker for undifferentiated spermatogonia stem cells) shows the specific and efficient PTBP1 deletion in Sertoli cells of *Ptbp1*^ΔSC^ mice. The arrowhead indicates mislocalized Sertoli cells in *Ptbp1*^ΔSC^ mice. Arrowheads indicate the PLZF-positive cells in which PTBP1 expression is intact in *Ptbp1*^ΔSC^ mice. (E-F) Hematoxylin and eosin (H&E)-stained testis and cauda epididymis sections from *Ptbp1*^fl/fl^ and *Ptbp1*^ΔSC^ mice at P35 and P100. Boxed regions were magnified in the middle panel. Arrows in F point to multinucleated giant cells in the seminiferous tubules of *Ptbp1*^ΔSC^ mice. Note very few spermatozoa are present in the cauda region.

We next assessed the depletion efficiency and specificity of PTBP1 in *Ptbp1*^ΔSC^ males by performing the triple immunofluorescence staining with antibodies against PTBP1, Sertoli cell marker GATA4, and undifferentiated spermatogonia cell marker PLZF. As shown in Figure 1D, PTBP1 was highly expressed in Sertoli cells in wild-type mice. Strong PTBP1 expression was also detected in PLZF-positive spermatogonia cells. In the *Ptbp1*^ΔSC^ mice, PTBP1 expression in Sertoli cells was diminished, while its expression in PLZF-positive cells remained intact (Figure 1D), indicating a specific and efficient PTBP1 deletion in Sertoli cells. Interestingly, some Sertoli cell nuclei were observed mispositioned in the seminiferous tubule lumen in the *Ptbp1*^ΔSC^ mice (Figure 1D).

Histological analysis indicated that severely affected *Ptbp1*^ΔSC^ mice had clusters of sloughed germ cells in the lumen but significantly less elongated spermatids. This was accompanied by a marked reduction of spermatozoa in the epididymis at P35 (Figure 1E). At P100, the defects became more severe. Elongated spermatids were barely detectable in *Ptbp1*^ΔSC^ mice (Figure 1F). Instead, multiple multinucleated giant cells appeared in the lumen (Figure 1F), and very few spermatozoa were present in the epididymis. This observation suggests that PTBP1 function in Sertoli cells is required for both the first and subsequent waves of spermatogenesis.

### Spermatid transformation is impaired in *Ptbp1*^ΔSC^ mice

Spermatogenesis includes several critical phases: mitosis, meiosis, spermiogenesis, and spermiation. We examined which step(s) of spermatogenesis was disrupted by Sertoli cell-specific PTBP1 deficiency. Production of undifferentiated spermatogonial cells was not affected in *Ptbp1*^ΔSC^ mice, as indicated by the comparable numbers of PLZF-positive cells in *Ptbp1*^fl/fl^ and *Ptbp1*^ΔSC^ mice (Supplementary Fig S1A and S1B). We next checked if PTBP1 deficiency affected meiosis by assessing germ cells positive for SYCP3 and γH2AX. SYCP3 is a DNA-binding protein and a component of the axial element of the synaptonemal complex, and its expression starts in leptotene germ cells and disappears in the late meiotic stage (43). γH2AX is required for the repair of DNA double-strand breaks in meiosis I and is localized in the nuclei of leptotene and zygotene spermatocytes and the sex chromosomes (formed “sex body”) of the pachytene spermatocytes (44). Both SYCP3-expressing and γH2AX-expressing germ cells were present in *Ptbp1*^ΔSC^ mice at P35 and P100, except that some labeled nuclei were mispositioned more toward the lumen at P100 (Supplementary Fig S1C to S1E). These observations suggest that PTBP1 in Sertoli cells is dispensable for the mitosis of spermatogonia and meiosis of spermatocytes, which prompted us to assess the defects in post-meiotic spermiogenesis.

During spermiogenesis in the mouse, haploid round spermatids undergo 16 steps of transformation to become elongated spermatids. Each step can be distinguished based on the changes in the acrosome cap and nuclear morphology of the younger generation of the spermatid (45,46). Staining of the acrosome using PAS and hematoxylin, spermiogenesis defects were assessed as previously reported (46). Transforming spermatids up to step 8 were present in *Ptbp1*^ΔSC^ mice at P35 and P100, although some were sloughed into the seminiferous lumen (Figures 2A and 2B). At P100, multinucleated giant cells containing spermatids formed in the seminiferous epithelium of *Ptbp1*^ΔSC^ mice and were undergoing apoptosis (Figure 2B, Supplementary Fig S2). TEM showed that a small number of spermatids in *Ptbp1*^ΔSC^ mice transformed beyond step 8 but had abnormal head morphologies and contained much larger portions of cytoplasm when compared to those in the control mice (Figure 2C and 2D). These observations indicate that spermiogenesis beyond step 8 was impaired in *Ptbp1*^ΔSC^ mice.

**Figure 2.**
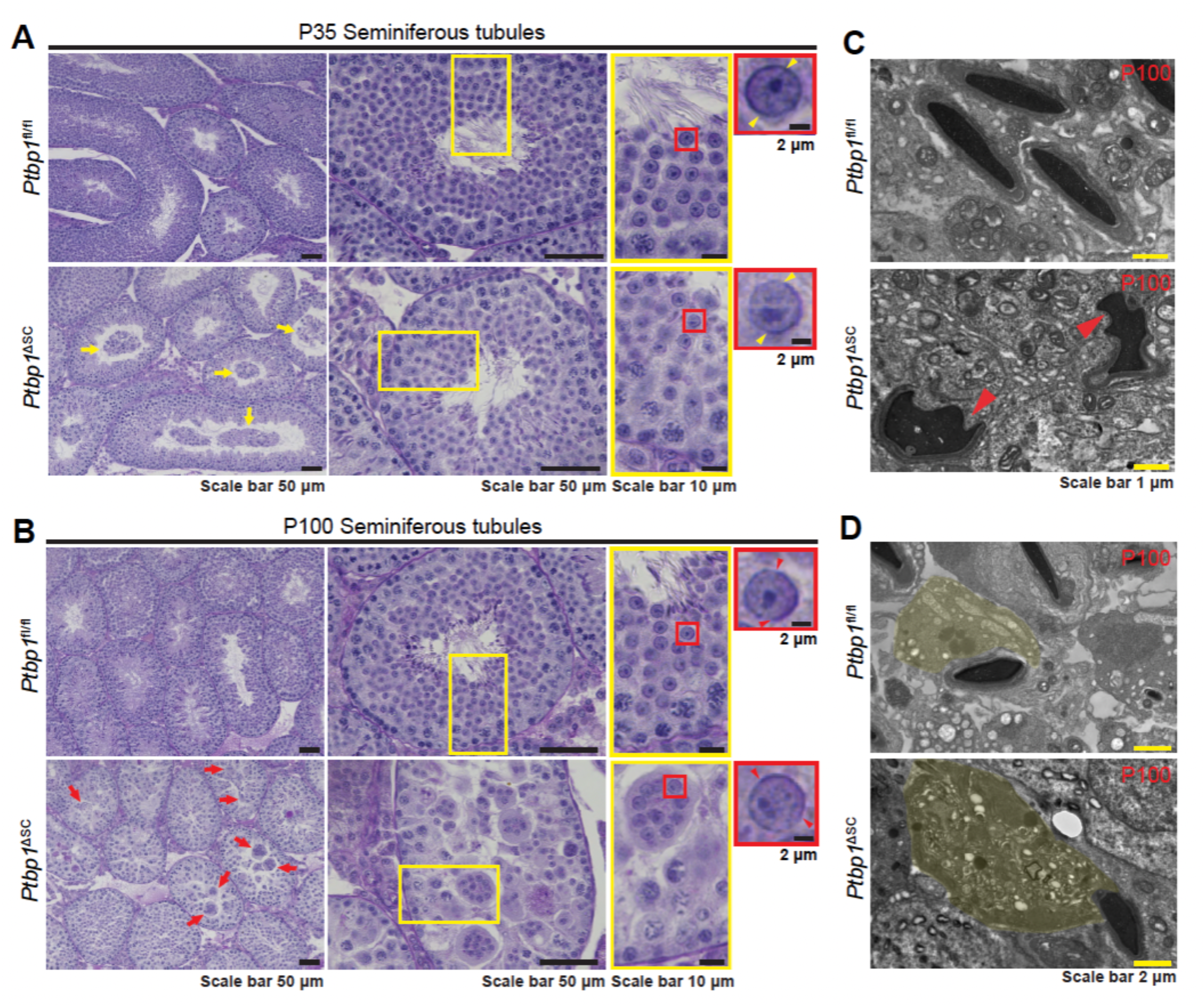
Spermiogenesis is arrested in *Ptbp1*^ΔSC^ mice. (A-B) The PAS and hematoxylin staining shows seminiferous cycles in the testes of *Ptbp1*^fl/fl^ and *Ptbp1*^ΔSC^ mice at P35 and P100. Regions in yellow boxes were magnified in the right panels. Yellow arrows in A point to sloughed germ cells at P35. Red arrows in B point to multinucleated giant cells at P100 in *Ptbp1*^ΔSC^ mice. Yellow arrowheads point to acrosome caps in step 8 spermatids. (C-D) Ultrastructural images of differentiating spermatids in *Ptbp1*^fl/fl^ and *Ptbp1*^ΔSC^ mice at P100. Red arrowheads in C point to the differentiating spermatids with abnormal head morphologies. Spermatid cytoplasm regions were pseudo-colored in yellow (D).

Taken together, these data indicate that *Ptbp1* functions in Sertoli cells to support spermiogenesis.

### Transcriptomic changes in the *Ptbp1*^ΔSC^ testis

To determine the molecular mechanism by which PTBP1 controls Sertoli cell function, we deep-sequenced poly(A) selected RNAs prepared freshly from age-matched wild-type and *Ptbp1*^ΔSC^ testes. We chose to assess the transcriptomic changes at P35 so that the observed changes represent the direct impact of Sertoli cell-specific PTBP1 deficiency on the first wave of spermatogenesis. PTBP1 deletion in Sertoli cells caused 501 altered splicing events within 401 genes and 140 differentially expressed genes (Figure 3A and 3B, Supplementary Table 4 and Table 5). Amongst them, 3 genes (*Gm28356*, *Porcn*, and *Il17re3*) exhibited significant differences in both mRNA abundance and splicing (Figure 3A).

**Figure 3.**
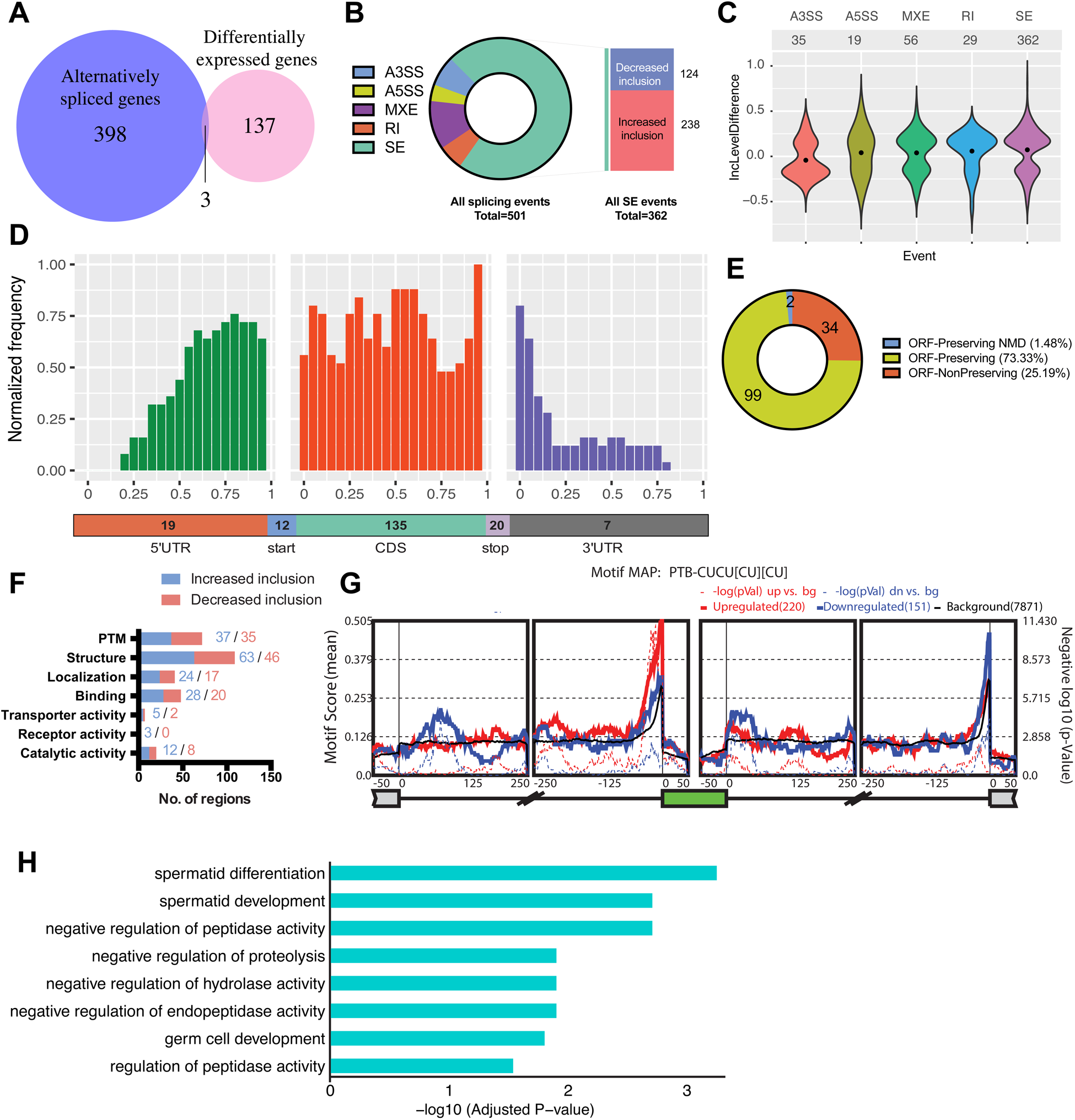
The transcriptome changes in the testis of *Ptbp1*^ΔSC^ mice. (A) A Venn diagram shows the number of genes that are differentially expressed or alternatively spliced in *Ptbp1*^ΔSC^ testes. Three genes(*Gm28356*, *Porcn*, and *Il17re3*) displayed changes in both gene expression and splicing. (B) Breakdown of 501 alternatively spliced events into various event categories. A3SS: alternative 3’ splice site, A5SS: Alternative 5’ splice site, MXE: Mutually exclusive exon, RI: Retained intron, SE: Skipped exon. (C) Violin plots demonstrate ΔPSI (Inclusion level difference) distributions of significantly altered splicing events in *Ptbp1*^ΔSC^ testes. (D) Metagene analysis of alternatively spliced exons detected in knockout mice by their positions on mRNA transcripts. Distribution along the transcript bins is shown on top, and the breakup of events into relative transcript regions is shown at the bottom. (E) Effect of alternatively spliced events induced by PTBP1 deficiency on the transcripts. Categories indicate if the exon inclusion preserves the open reading frame (ORF) and if the inclusion transcript is subjected to Non-sense Mediated Decay (NMD). (F) Exon ontology-based distribution of skipped exons, either increase or decrease in inclusion, based on encoded protein features. PTM: post-translational modification. (G) shows the relative enrichment of [CT]-rich motif near cassette exons (represented in green) that displayed a significant increase (red curve) or decrease (blue curve) in inclusion in *Ptbp1*^ΔSC^ testes. Alternatively spliced exons were identified using rMATS, and a motif map was constructed using RMAPs (with a 50-nucleotide sliding window). The set of background cassette exons is represented in black. (H) Gene ontology analysis demonstrates the top 8 biological processes enriched among differentially expressed genes.

The majority of splicing changes in the PTBP1-deficient testis were exon skipping events (72%), but changes in alternative 5’ or 3’ splice sites, intron retention, and mutually exclusive exons were also detected (Figure 3B). Notably, most affected exons displayed increased inclusion in the PTBP1-deficient testis (Figure 3B and 3C), which is consistent with the previous findings from us and others that PTBP1 primarily functions as a repressor of splicing (47,48). We next investigated the spatial distribution of alternatively spliced exons along their associated transcripts, as previously described (49). Metagene analysis revealed that approximately 70% of differentially spliced exons were located within coding sequences (CDS), and a sizeable number of those (17%) encoded alternate START or STOP codons (Figure 3D).

The CDS-mapped PTBP1-regulated exons were further classified based on whether they were open reading frame preserving (exon length is a multiple of 3). We found that 75% of the PTBP1-regulated exons preserve the open reading frame, of which only 1.5% were predicted to undergo nonsense-mediated RNA decay (Figure 3E). This result is consistent with our transcriptome data wherein the majority of the mRNAs harboring PTBP1-regulated exons in the testis do not exhibit a significant change in their overall abundance (Figure 3A). On the contrary, these exons are likely to alter the intrinsic structure and function of the encoded proteins. To further probe the functional properties and features of these exons, we performed exon ontology analysis, which revealed significant enrichment for sequences encoding molecular regions for the structure, post-translational modifications, cellular localization, binding, and transporter, receptor, or catalytic activities (Figure 3F).

We found a substantial overrepresentation of the CUCUCUCU motif near the 3’ splice site of exons that are more included upon PTBP1 depletion (Figure 3G). These CU/pyrimidine-rich motifs represent direct binding motifs for PTBP1 (50–52), suggesting that in the testis, PTBP1 suppresses the inclusion of many alternate exons by directly binding to this motif in the upstream introns.

Gene ontology analysis of differentially expressed genes revealed that the top biological process affected is spermatid differentiation (Figure 3H), which is consistent with our histological observation of defective spermiogenesis in *Ptbp1*^ΔSC^ mice (Figure 2A and 2B).

### Loss of PTBP1 altered the splicing of actin cytoskeleton regulators

Since the *Ptbp1* gene was specifically deleted in Sertoli cells in *Ptbp1*^ΔSC^ mice, the transcriptomic changes in Sertoli cells represent the direct consequence of PTBP1 deficiency. We thus compared our data with published Sertoli cell-expressing genes (53,54), and identified 27 Sertoli cell-enriched genes that are alternatively spliced upon PTBP1 deficiency (Supplementary Table 6). Strikingly, 8 of them (*Espn*, *Tnik*, *Gm14569*, *Ppp1r9a*, *Appl2*, *Rab11fip3*, *Phactr2*, and *Camkk2*) were reported to be functionally required for actin cytoskeleton organization (18,55–64). Except for *Espn*, splicing of more than one exon of each gene was affected by PTBP1 deletion, either in exon skipping or mutually exclusive AS (Figure 4A). Most affected exons had increased inclusion in *Ptbp1*^ΔSC^ mice (Figure 4), indicating that PTBP1 acts predominantly as a repressor of exon inclusion in these genes. Using the PCR-based splicing assay, we validated 7 splicing events in 7 genes, including 5 actin regulators (Figure 4B and 4C, Supplementary Fig S3).

**Figure 4.**
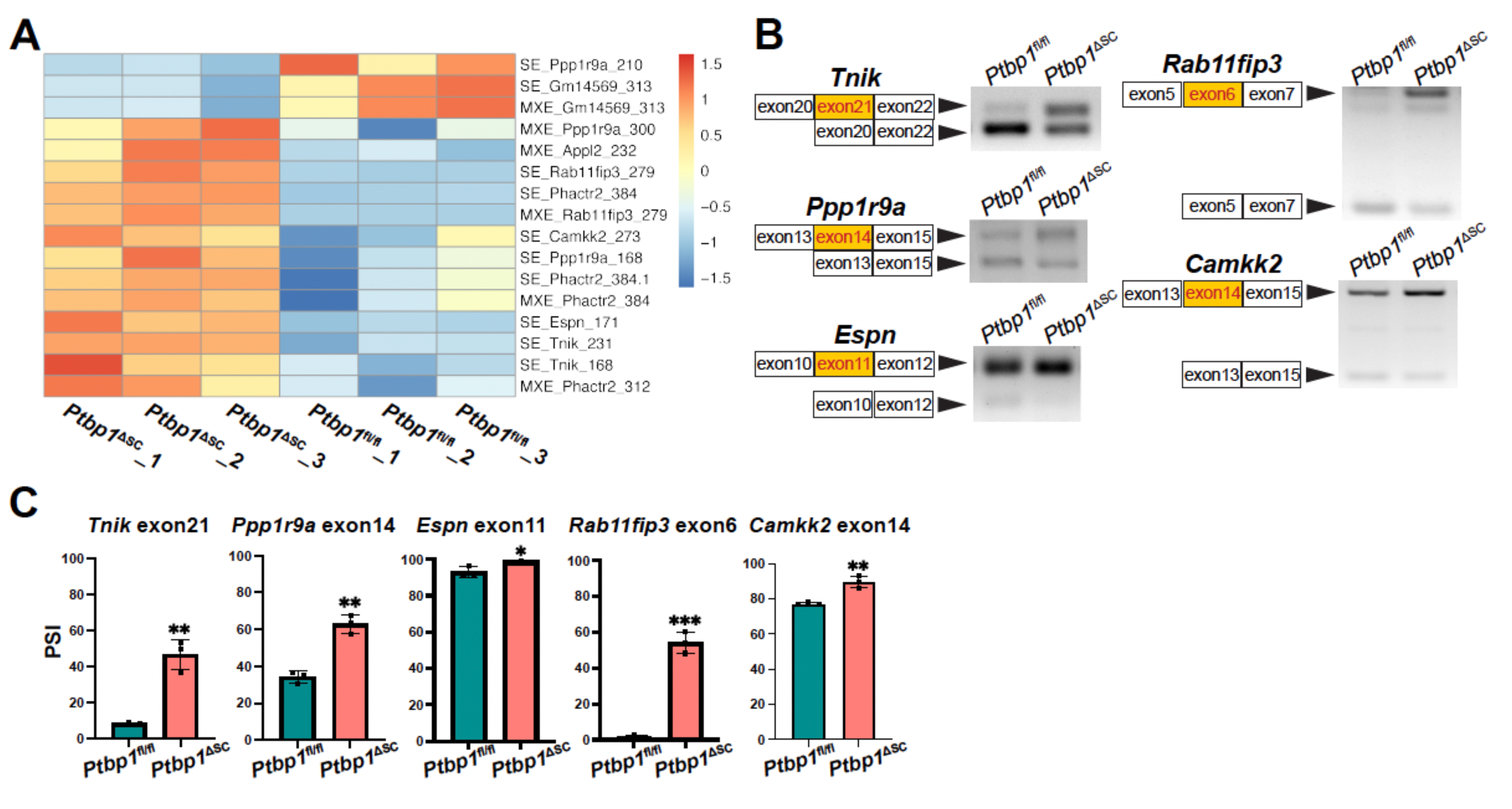
PTBP1 deficiency induced aberrant splicing of Sertoli cell-enriched actin regulators. (A) The heatmap shows the PSI values of exons in 8 actin regulators enriched in Sertoli cells. (B-C) PCR-based splicing assays validate 5 alternatively spliced events in 5 actin regulatory genes. Representative gel images are shown in B, and the quantifications of PSI are shown in C. 3 *Ptbp1*^ΔSC^ mice and 3 sibling littermate *Ptbp1*^fl/fl^ mice were assessed. Data are presented as mean± SD, n=3. *P<0.05, **P<0.01, ***P<0.001.

Notably, none of the aberrantly spliced Sertoli cell genes has been reported to be involved in Sertoli cell maturation. To confirm Sertoli cell maturation was not affected by PTBP1 deficiency, we checked the expression of androgen receptor, a marker of Sertoli cell maturation (35). Indeed, *Ptbp1* deletion did not affect the androgen receptor expression in Sertoli cells (Supplementary Fig S4), suggesting that PTBP1-mediated splicing control is dispensable for Sertoli cell maturation.

### Loss of PTBP1 disrupted the BTB function and actin cytoskeleton organization

Given that PTBP1 deficiency altered the splicing of 8 actin regulators, we examined if F-actin organization was impaired in *Ptbp1*^ΔSC^ Sertoli cells. In the control mice, thick F-actin bundles were organized at the basal ES near the basement membrane and apical ES surrounding the heads of transforming spermatids (Figure 5A). In contrast, the F-actin organization was completely disrupted in *Ptbp1*^ΔSC^ testes. Tangled and aggregated actin bundles were randomly distributed in the lumen of the seminiferous tubule (Figure 5A). At the basal region, F-actin bundles were noticeably thinner when compared to those of the control mice (Figure 5A, red inset). Elongated spermatids were rarely detected in *Ptbp1*^ΔSC^ mice. Occasionally, a thin layer of F-actin bundle was detected around the step 8 round spermatids in the multinucleated giant cells (Figure 5A, yellow inset). Since F-actin is a crucial component of ES, we next examined the ES structure in *Ptbp1*^ΔSC^ mice. We assessed the localization of ESPN, an ES marker (18,55). Indeed, ESPN was mislocalized at both the apical and basal regions of Sertoli cells (Figure 5B), suggesting impaired ESs.

**Figure 5.**
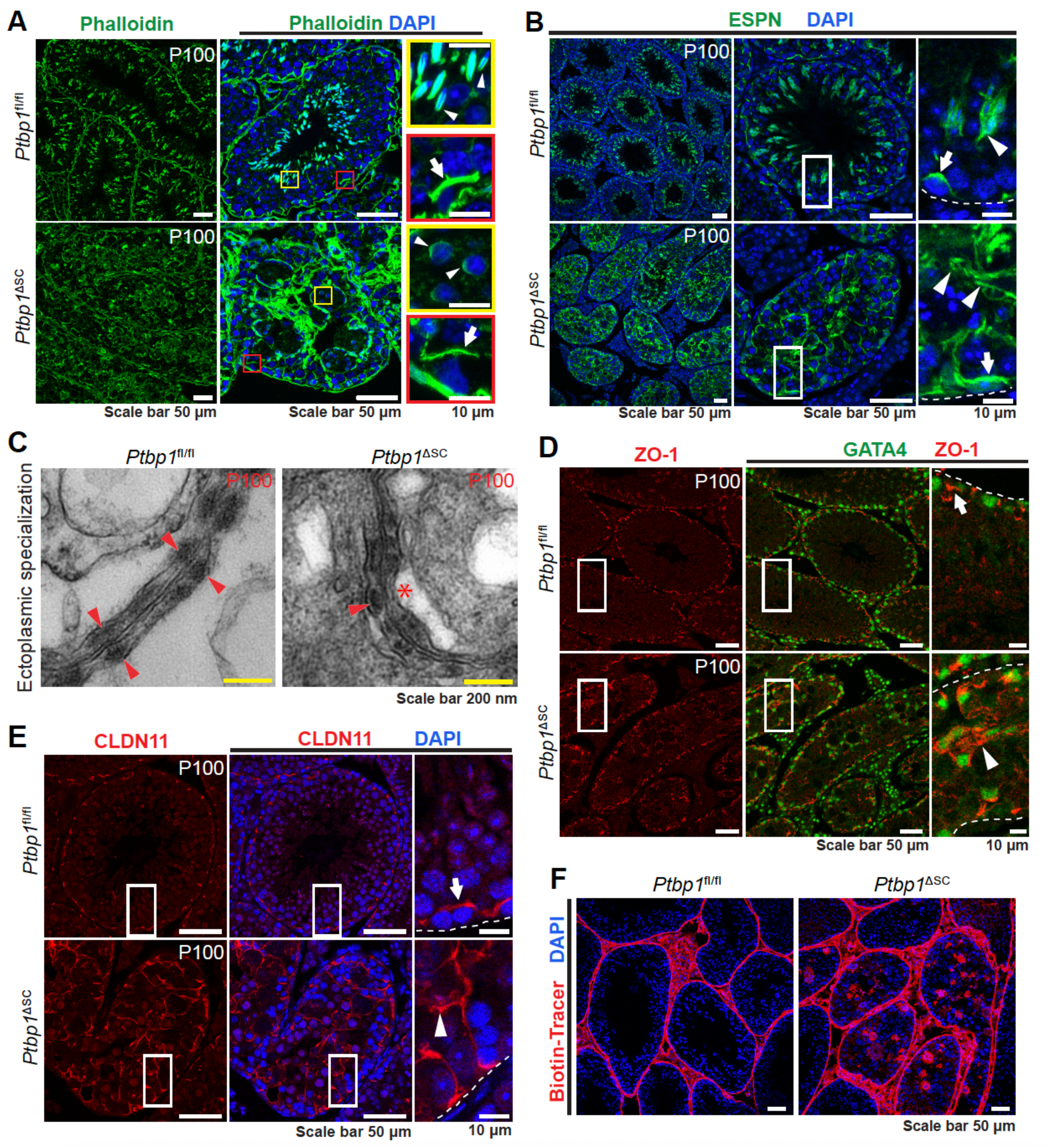
*Ptbp1*^ΔSC^ mice had disorganized F-actin at the ES and impaired BTB integrity. (A) Phalloidin staining shows F-actin organization. Apical ES ( yellow boxes) and basal ES (red boxes) are magnified in insets. Note that Sertoli cells of *Ptbp1*^ΔSC^ mice exhibit abnormal actin assembly at basal ES (arrows) and apical ES (arrowheads). (B) Immunofluorescence staining using the antibody against ESPN shows impaired basal ES (arrows) and apical ES (arrowheads). (C) TEM images show the basal ES. Arrowheads indicate the packed F-actin bundles in the ES of adjoining Sertoli cells. The asterisk points to the region where F-actin bundles are missing in the Sertoli cells of *Ptbp1*^ΔSC^ mice. (D and E) Immunofluorescence staining using the antibodies against GATA4, ZO-1, or CLDN11 shows the distribution of tight junction proteins. Boxed regions are magnified on the right. Arrows show the normal distribution of ZO-1 and CLDN11 at the basal region of Sertoli cells in *Ptbp1*^fl/fl^ mice. Arrowheads show the mislocalized ZO-1 and CLDN11 in *Ptbp1*^ΔSC^ mice. (**F**) Sulfo-NHSLC-biotin staining shows impaired BTB integrity.

To further assess the ES structure at the ultrastructural level, we performed the TEM. In the control mice, the basal ESs were formed at the neighboring Sertoli cells, structurally characterized by a layer of actin bundles sandwiched between a cistern of the endoplasmic reticulum and the nearby plasma membrane of the Sertoli cell (Figure 5C). The F-actin bundles were symmetrically assembled in the basal ESs of the adjoining Sertoli cells (Figure 5C, left panel). On the contrary, F-actin bundles in the PTBP1-deficient Sertoli cells appeared asymmetrical and irregular (Figure 5C, right panel). Since the basal ES connects other junctions to form BTB, we checked how PTBP1 deficiency affected the distribution of other junction proteins. As shown in Figure 5D and 5E, the tight junction proteins ZO-1 and CLAUDIN-11 (CLDN11) were no longer restricted between neighboring Sertoli cells near the basement membrane in *Ptbp1*^ΔSC^ mice. Instead, they diffused into the apical region (Figure 5D and 5E). We examined the functional integrity of BTB by assessing its permeability using a small molecule, sulfo-NHSLC-biotin, as previously described (36). Indeed, the BTB permeability in PTBP1-deficient Sertoli cells was significantly increased, as indicated by the diffusion of sulfo-NHSLC-biotin to the adluminal compartment (Figure 5F).

Together, our findings indicate that loss of PTBP1 in Sertoli cells disrupted the organization of F-actin cytoskeleton at the apical and basal ES and impaired the BTB function.

### PTBP1 represses the inclusion of exon 14 of *Tnik* in Sertoli cells

We next investigated how PTBP1-mediated AS controls the organization of actin cytoskeleton in Sertoli cells. We focused on the impact of AS of exon 14 of *Tnik* for three reasons. First, its human counterpart exon is alternatively spliced in the human testis, but its physiological significance has not been studied (65). Secondly, the inclusion of this exon in human *Tnik* inhibits F-actin assembly in neuronal cells *in vitro* (65). Thirdly, Human and mouse *TNIK* share a 99% protein similarity(58), and mouse and human exon 14 encode 100% identical amino acids, suggesting TNIK may play a conserved role in the two species (Supplementary Fig S5). Notably, this exon in human *Tnik* has been described as exon 15 previously (65,66). However, based on the most recent genome sequence information from the UCSC genome and the National Center for Biotechnology Information (NCBI), this exon is the 14th of the 33 exons of human *Tnik*. As such, we named it human exon 14 in this report.

*Tnik* encodes a germinal center kinase family member that interacts with Traf2 and NCK(67). *TNIK* protein comprises an N-terminal kinase domain, a C-terminal regulatory domain, and an intermediate region (58,68). *Tnik* exon 14 encodes 29 amino acids in the intermediate region (Supplemental Fig S5). Interestingly, under homeostatic conditions, AS of *Tnik* exon 14 in the mouse testis was developmental stage specific. *Tnik* Exon 14 was predominantly included at P8 but was mostly excluded at P14 (Figure 6A and 6B). At P35 and P120, the inclusion of exon 14 became predominant again. Depletion of PTBP1 resulted in a strong inclusion of exon 14 in *Tnik* in all stages examined, suggesting that PTBP1 functions as a repressor of exon 14 inclusion. Using a commercial antibody that recognizes all TNIK protein isoforms, we assessed the total protein level of TNIK in the testes of P9 *Ptbp1*^ΔSC^ mice and their sibling control mice by Western blotting. No significant changes in the TNIK protein amount were found in *Ptbp1*^ΔSC^ mice (Supplementary Fig S6), suggesting that exon 14 inclusion induced by PTBP1 deficiency did not induce nonsense-mediated RNA decay of *Tnik* transcripts.

**Figure 6.**
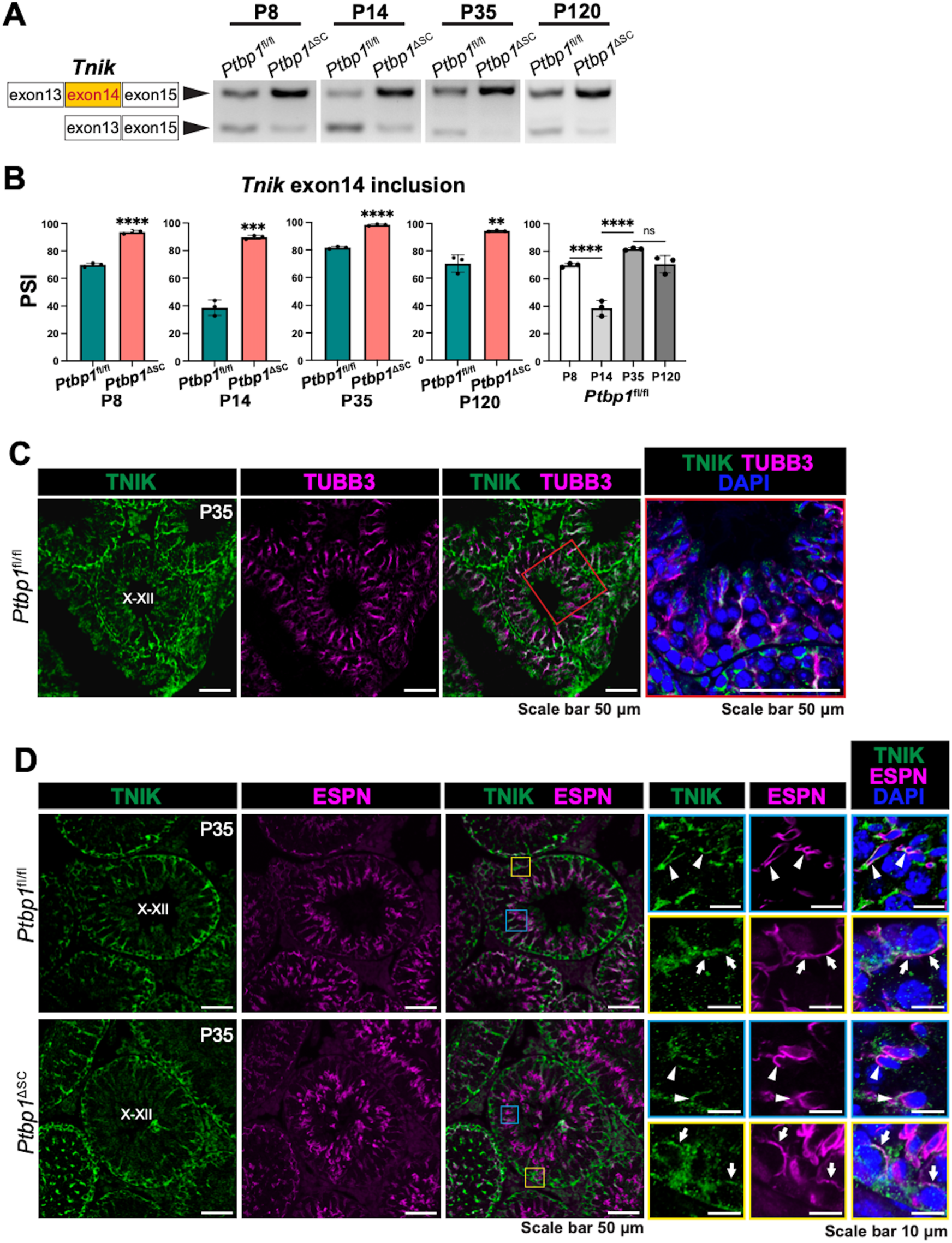
PTBP1 deficiency increases the inclusion of *Tnik* exon 14 and alters TNIK protein distribution. (A-B) PCR-based splicing assays demonstrate the splicing patterns of *Tnik* exon 14 at different ages. Representative gel images in A show the presence of exon 14-included and -excluded *Tnik* isoforms in the testes at different ages. Quantifications of PSI are shown in B. 3 *Ptbp1*^ΔSC^ mice and 3 sibling littermate *Ptbp1*^fl/fl^ mice were assessed at each age. Data are presented as mean± SD, n=3. **P<0.01, ***P<0.001, ****P<0.0001. ns, not significant. (C) Double immunofluorescence staining using antibodies against TNIK and TUBB3 shows TNIK expression in the Sertoli cell cytoplasm. Boxed regions were magnified in the right panels. (D) Double immunofluorescence staining using antibodies against TNIK and ESPN shows TNIK expression in the apical ES (blue boxes) and basal ES (yellow boxes). Boxed regions are magnified in insets. PTBP1 deficiency altered TNIK distribution at both the apical ES (arrowheads) and basal ES (arrows).

We next checked TNIK cellular localization in Sertoli cells by performing double immunostaining with antibodies against TNIK and TUBB3 (also known as Class III β-TUBULIN). TUBB3 was used as a Sertoli cell cytoplasm marker (69). TNIK was highly accumulated at the basal region of Sertoli cells near the basement membrane. Its expression further radiated to the apical side, parallel to the Sertoli cell’s long axis, and then became branched near the apical lumen (Figure 6C). We next performed double immunostaining with antibodies against TNIK and ESPN to examine its presence at the basal and apical ES and the impact of PTBP1 deficiency on its distribution. In the control mice, TNIK was accumulated at the apical ES surrounding the head of the transforming spermatid as well as at the basal ES, partially overlapping with ESPN (Figure 6D, insets). Deletion of PTBP1 caused the sparse distribution of TNIK at both apical and basal ES, but its accumulation along the long axis of Sertoli cells was not affected (Figure 6D).

### *Tnik* exon 14 exclusion promotes F-actin assembly in Sertoli cells

The inclusion of human *Tnik* exon14 has been found to inhibit F-actin bundle formation in cultured neuronal cells (65). This promoted us to examine the function of mouse *Tnik* exon 14 in regulating Sertoli cell actin cytoskeleton organization. To this end, we cultured Sertoli cells derived from *Ptbp1*^ΔSC^ and *Ptbp1*^fl/fl^ mice *in vitro*. F-actin bundles were noticeably thinner in PTBP1-deficient Sertoli cells than the control Sertoli cells (Figure 7A). To determine if defective F-actin bundle assembly in PTBP1-deficient Sertoli cells was caused by exon 14 inclusion in *Tnik*, we treated *Ptbp1*^ΔSC^ Sertoli cells with splicing-inhibiting ASO to block the inclusion of exon14 in *Tnik* (Fig 7B to 7D). Remarkably, blocking exon14 inclusion partially restored the F-actin bundle assembly in PTBP1-deficient Sertoli cells (Fig 7E). This finding demonstrates that PTBP1 represses *Tnik* exon14 inclusion to promote F-actin bundle formation in Sertoli cells.

**Figure 7.**
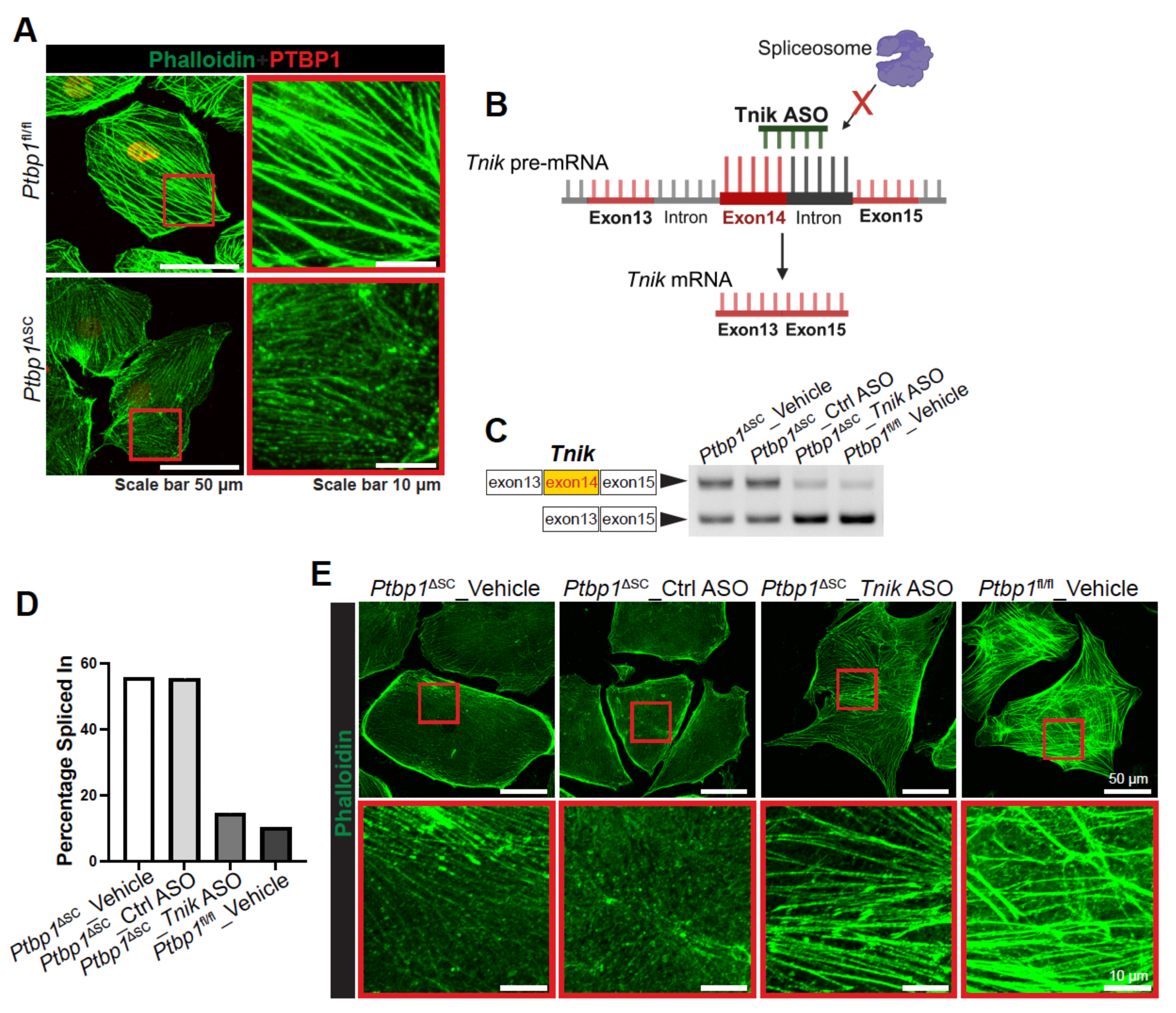
PTBP1 represses the inclusion of *Tnik* exon14 to promote F-actin assembly in Sertoli cells. (A) Double fluorescence staining of PTBP1 and F-actin shows reduced F-actin bundle formation in PTBP1-deficient Sertoli cells cultured *in vitro*. Boxed regions are magnified in the right panels. (B) A schematic diagram shows the action of *Tnik* splicing-inhibiting ASO in blocking *Tnik* exon 14 inclusion. *Tnik* ASO binds to the 3’end of *Tnik* exon 14 and the adjacent 3’ splice site in the intron region, which prevents the recognition of 3’ splice site by the spliceosome. This leads to the skipping of exon 14. (C and D) PCR-based splicing assays show the *Tnik* ASO treatment reduced the inclusion of *Tnik* exon 14 in PTBP1-deficient Sertoli cells. The data shown are representative of three independent experiments. (E) Phalloidin staining shows F-actin bundles in *Tnik* ASO- or vehicle-treated Sertoli cells derived from *Ptbp1*^fl/fl^ and *Ptbp1*^ΔSC^ mice. Sertoli cells from *Ptbp1*^fl/fl^ were used as a reference for normal F-actin bundle formation. Boxed regions are magnified in the bottom panels. Note F-actin bundles in *Tnik* ASO-treated PTBP1-deficient Sertoli cells are much thicker than those in vehicle-treated ones.

Collectively, our results reveal a novel mechanism wherein PTBP1-mediated AS of Sertoli cell genes controls the actin cytoskeleton dynamics. This splicing control organizes the F-actin cytoskeleton at the apical ES for spermiogenesis and at the basal ES for the BTB function. Mechanistically, it is executed at least in part through repressing the inclusion of *Tnik* exon 14 to promote actin bundle formation (Figure 8)

**Figure 8.**
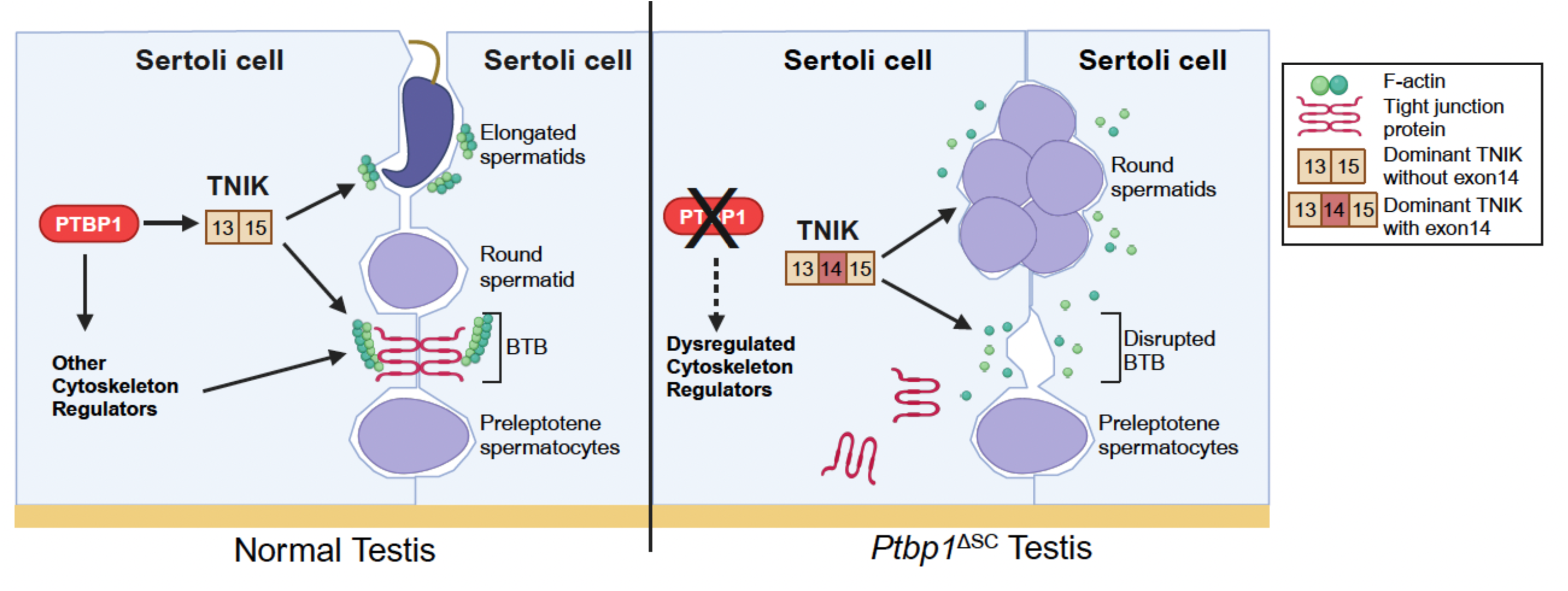
A proposed model of PTBP1 in regulating actin cytoskeleton organization in Sertoli cells. PTBP1 controls AS of actin cytoskeleton regulators in Sertoli cells to support the apical and basal ES function, which is essential for BTB integrity, germ cell transport, adhesion, and differentiation. Of particular, PTBP1 represses the inclusion of *Tnik* exon 14 to promote the F-actin bundle formation.

## DISCUSSION

Sertoli cells support spermatogenesis through a wide variety of functions, ranging from providing nutrients and hormones to germ cells to structurally supporting their translocation, positioning, differentiation, and release into the lumen of the seminiferous tubule (2). Many of these functions rely on the remarkably dynamic actin cytoskeleton in Sertoli cells. F-actin is concentrated at the apical and basal ES of Sertoli cells, where specialized adhesion junctional complexes enable Sertoli-Sertoli cell and Sertoli-germ cell interactions. The basal ESs of neighboring Sertoli cells are associated with other junctions to form a structural component of the BTB and protect immunogenic meiotic and post-meiotic germ cells from autoimmune attacks. The apical ES of the Sertoli cell forms around the head of the differentiating spermatid to facilitate the latter steps of spermiogenesis. Both apical and basal ESs undergo cyclic breakdown and reconstruction to orchestrate spermatogenesis. Timely controlled actin bundle formation and disassembly in the ES is key to the proper ES turnover.

In this report, we explored the post-transcriptional regulatory mechanism that controls actin cytoskeleton dynamics by conducting genome-wide profiling of PTBP1-mediated AS in Sertoli cells. We show that PTBP1-mediated AS in Sertoli cells controls actin cytoskeleton organization in ESs,. Mechanistically, PTBP1 controls AS of 8 genes involved in actin cytoskeleton assembly. Of particular note, PTBP1 represses the inclusion of *Tnik* exon 14 in Sertoli cells to promote actin bundling. TNIK is a Serine/threonine kinase required for synapse composition, activity, and dendritic growth in neuronal cells (65,70). Overexpression of human exon 14-included but not excluded *Tnik* in neuronal cells reduced F-actin density in the growth cone of neurites and impaired neuronal development *in vitro* (65), suggesting that exon 14 is required for *Tnik*-mediated inhibition of actin bundle formation. In this study, we used a knockout mouse model to show that PTBP1 is an important splicing regulator of *Tnik* exon 14 in Sertoli cells *in vivo*. Further, using the splicing-inhibiting ASO that blocks the exon 14 inclusion, we provide direct evidence demonstrating that the exclusion of this exon in *Tnik* attenuates the inhibitory activity of *Tnik* in actin bundling in Sertoli cells. Our results thus reveal a conserved role of this exon in regulating TNIK function in mice. Notably, *Tnik* exon 14 is also alternatively spliced in human testes (65). Compared to human testes, wherein the exon 14-excluded *Tnik* isoform is predominant (65), the mouse testis contains primarily the exon 14-included *Tnik* isoform (Figure 6A). This discrepancy is likely caused by the fact that the human testis RNA sample used for assessing the *Tnik* exon 14 splicing was from a mixture of testes collected from individuals of different ages(65), which potentially masks the age-specific splicing difference of *Tnik*.

It is worth noting that while how the exclusion of exon 14 attenuates the inhibitory activity of TNIK on actin polymerization is not fully understood yet, evidence in the literature suggests that this mechanism is involved in neurite development, a biological process highly dependent on actin dynamics. *Tnik* exon 14 encodes 29 amino acids in the intermediate region of TNIK. This region of TNIK has been reported to mediate TNIK interaction with an E3 ubiquitin ligase NEDD4-1. The exon 14-included *Tnik* transcript produces a TNIK isoform capable of forming a complex with NEDD4-1 and promoting the turnover of small GTPase RAP2A, which controls the neurite development(70). It would be of great interest to determine if AS of *Tnik* exon 14 regulates actin bindle assembly via controlling the ubiquitin/proteasome-dependent turnover of RAP2A.

We found that both exon 14-included and excluded *Tnik* isoforms were present in mouse Sertoli cells, and their ratio fluctuated during the testis development (Figure 6A). While further studies are needed to determine the physiological importance of such fluctuation, it is tempting to speculate that the ratio of *Tnik* splicing isoforms may be timely controlled to coordinate actin dynamics during an epithelial cycle. During this process, PTBP1 may work with other RNA-binding proteins to titrate the ratio of *Tnik* isoforms. Interestingly, RNA-binding proteins NOVA-1 and TDP-43 compete to control the skipping of human *Tnik* exon 14 in HEK293T cells(65). TDP-43 is expressed in Sertoli cells, and its deficiency in Sertoli cells disrupts spermatogenesis and results in testicular defects similar to what we observed in the *Ptbp1*^ΔSC^ mice, including impairment of the BTB function and mislocalized germ cells (71). It would be interesting to determine if TDP-43 or other RNA-binding proteins work with PTBP1 in Sertoli cells to control the splicing of *Tnik* exon 14.

Besides exon 14, human *Tnik* contains two other alternative exons, exon 16 and exon 21, which encode 53 and 7 amino acids in the intermediate region, respectively. All three exons are alternatively spliced to generate 8 protein isoforms of human TNIK (68). We found that loss of PTBP1 in Sertoli cells increased the inclusion of both *Tnik* exon 14 and 21 (Fig 4A). In this study, we focused studying the role of exon 14. Whether exon 21 of *Tnik* plays a role in regulating actin cytoskeleton organization needs further study.

In addition to TNIK, PTBP1 regulates the AS of 7 other actin regulators in Sertoli cells. Other than ESPN, a well-known ES-localized actin-binding protein important for actin bundling(18,55), their functions in regulating actin cytoskeleton organization in Sertoli cells have not been studied. Gm14569 is an ortholog of human KIAA1210. It is localized at the ES and has been reported to interact with tight junction proteins such as ZO1(56), but its role in regulating actin assembly in the ES is unknown. CAMKK2 is a kinase that regulates actin assembly to guide cell adhesion, migration, and morphogenesis during tissue homeostasis and cancer metastasis (57–60). PPP1R9A (also known as neural-specific F-actin-binding protein, NEURABIN1) is a kinase that binds to F-actin to promote neurite formation and synaptic transmission(61–63). PHACTR2 (Phosphatase and actin regulator 2) is highly enriched in neurons and has a strong actin-binding activity (64). While our findings reveal an important role of aberrantly spliced *Tnik* in causing defective actin cytoskeleton organization, we cannot exclude the possibility that mis-splicing of the remaining 7 genes provoked by PTBP1 deficiency also contributes to these defects. Follow-up studies are required to determine the contribution of these genes in regulating Sertoli cell functions.

## DATA AVAILABILITY

All raw RNA-seq data files are available for download from the Gene Expression Omnibus (accession number GSE267924).

## ACKNOWLEDGMENTS

We thank Ms. Cate Wallace and Dr. Jade Wang for the technical support of TEM. We thank Dr. Waqar Arif for sharing the computational tools. The rat anti-ZO1 antibody was obtained from the Developmental Studies Hybridoma Bank, which is created by the NICHD of the NIH and maintained at the University of Iowa, Department of Biology, Iowa City, IA 52242.

## AUTHOR CONTRIBUTIONS

Y.W. designed the project, performed the experiments, interpreted the data, and drafted the manuscript. V.C.U. and S.C. performed computational analysis and interpreted the data, D.Y. performed the experiments. J.J., Y-J.W., K.L.N, and P.L. performed data analysis. A.R., R.H. and H.Q. interpreted the result and intellectually contributed to the experimental design, K.K., J.Y., and A.K. designed the research and supervised the study. W.M. designed the research, performed the experiments, interpreted the data, supervised the study, and revised the manuscript.

## Supplementary figure legends

**Supplementary Fig 1.**
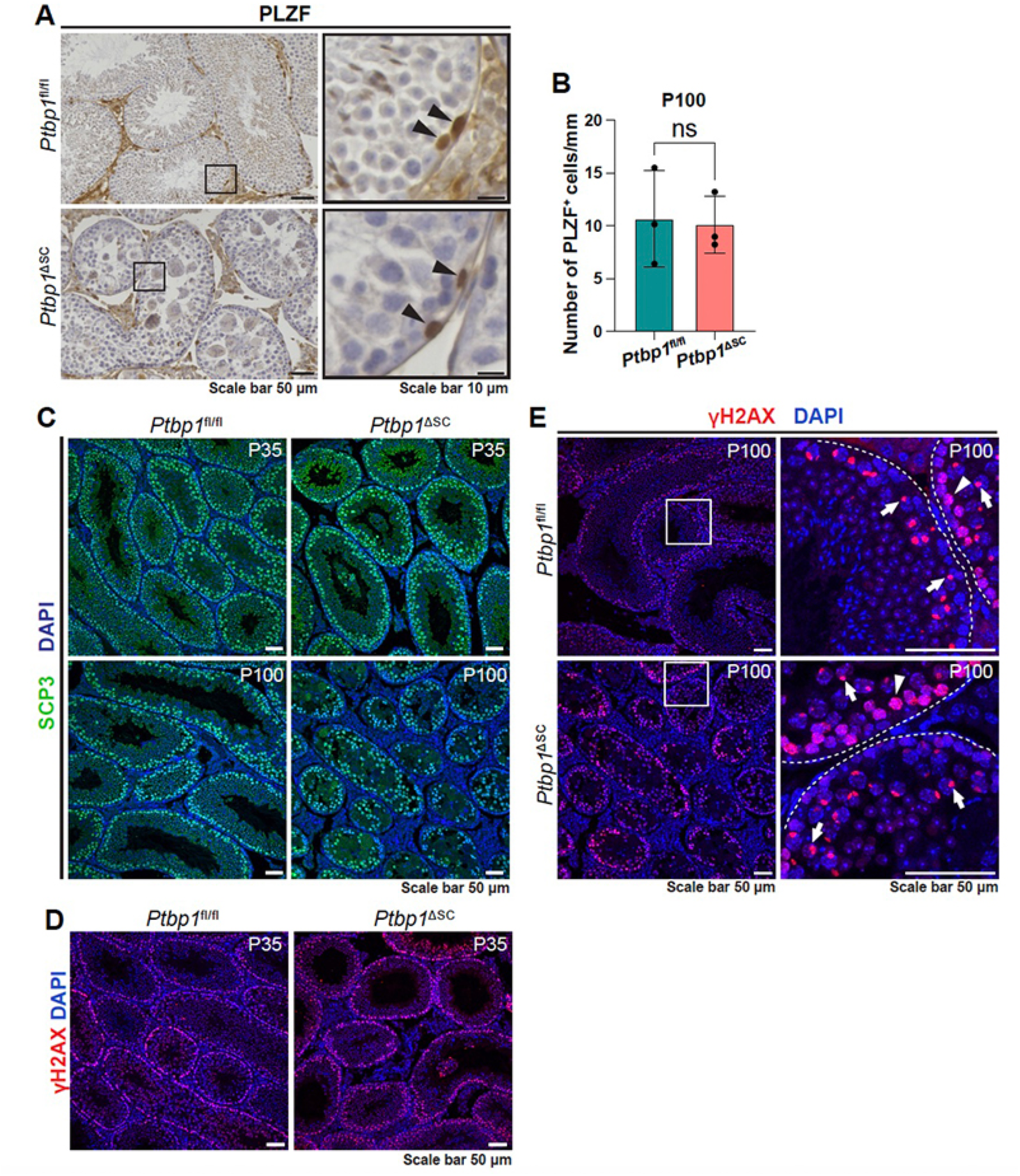
Germ cell mitosis and meiosis were not affected by *Ptbp1* deletion. **(A)** Immunohistochemistry staining with an antibody against PLZF shows undifferentiated spermatogonia cells in the testes of *Ptbp1*^fl/fl^ and *Ptbp1*^ΔSC^ mice. Boxed regions are magnified on the right. Arrowheads indicate PLZF-positive spermatogonia cells. (**B**) Quantification of PLZF-positive spermatogonia cells. 329 seminiferous tubules from the testis sections of 3 *Ptbp1*^fl/fl^ mice and 390 tubules from 3 *Ptbp1*^ΔSC^ mice were assessed. The number of PLZF-positive cells in each tubule was divided by the perimeter of the tubule (mm). The resulting values were averaged in each mouse and compared between the two groups. NS, not significant. (C-E) Immunofluorescent staining shows SYCP3-positive spermatocytes or γH2AX-positive germ cells at the leptotene and zygotene stage (arrowheads) in *Ptbp1*^fl/fl^ and *Ptbp1*^ΔSC^ mice at P35 and P100. Boxed regions were magnified in the right panels in E. γH2AX is condensed in the sex body of germ cells at the pachytene stage(arrows). Dashed lines indicate the edges of seminiferous tubules.

**Supplementary Fig 2.**
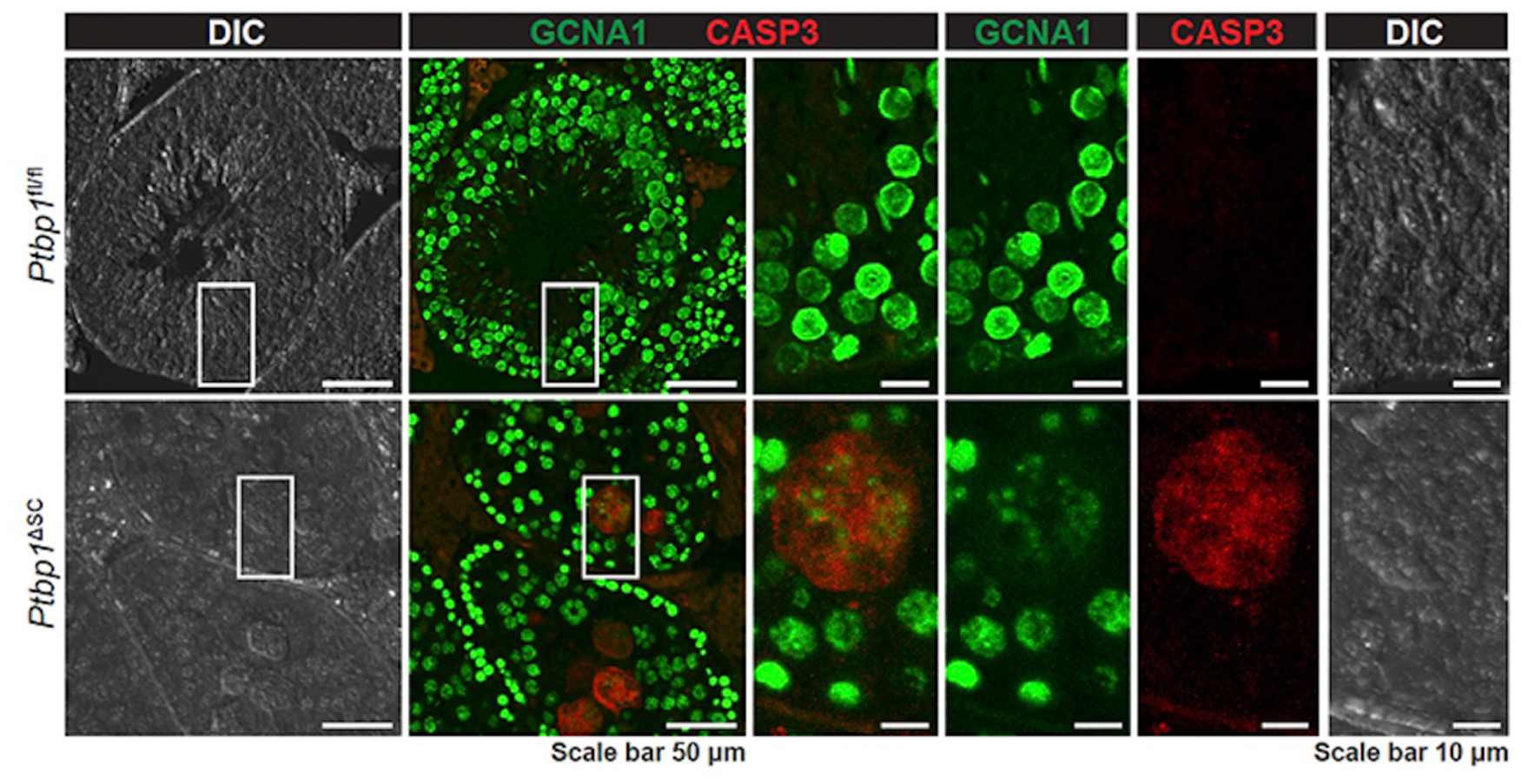
Germ cells formed apoptotic multinucleated giant cells in *Ptbp1*^ΔSC^ mice. Double immunofluorescence staining using the antibodies against GCNA1 and cleaved CASPASE3 (CASP3) shows the multinucleated giant cells formed by germ cells underwent apoptosis. GCNA1 is used for labeling germ cells. Boxed regions are magnified in the right panels.

**Supplementary Fig 3.**
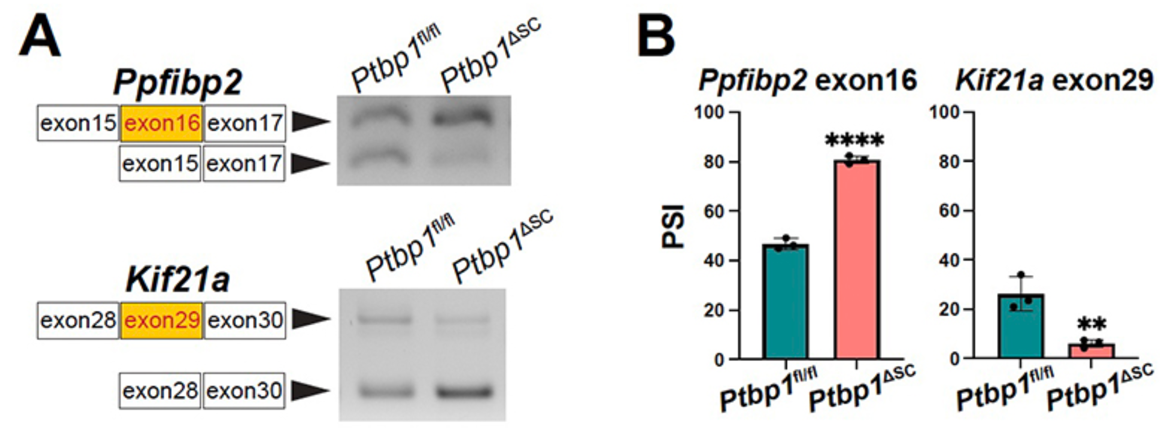
Validation of alternatively spliced events in *Ptbp1*^ΔSC^ mice. (A) Representative gel images show alternatively spliced events in *Ptbp1*^ΔSC^ mice. (B) shows the quantifications of PSI. 3 *Ptbp1*^ΔSC^ mice and 3 sibling littermate *Ptbp1*^fl/fl^ mice were assessed. Data are presented as mean± SD. n=3. **P<0.01, ****P<0.0001.

**Supplementary Fig4.**
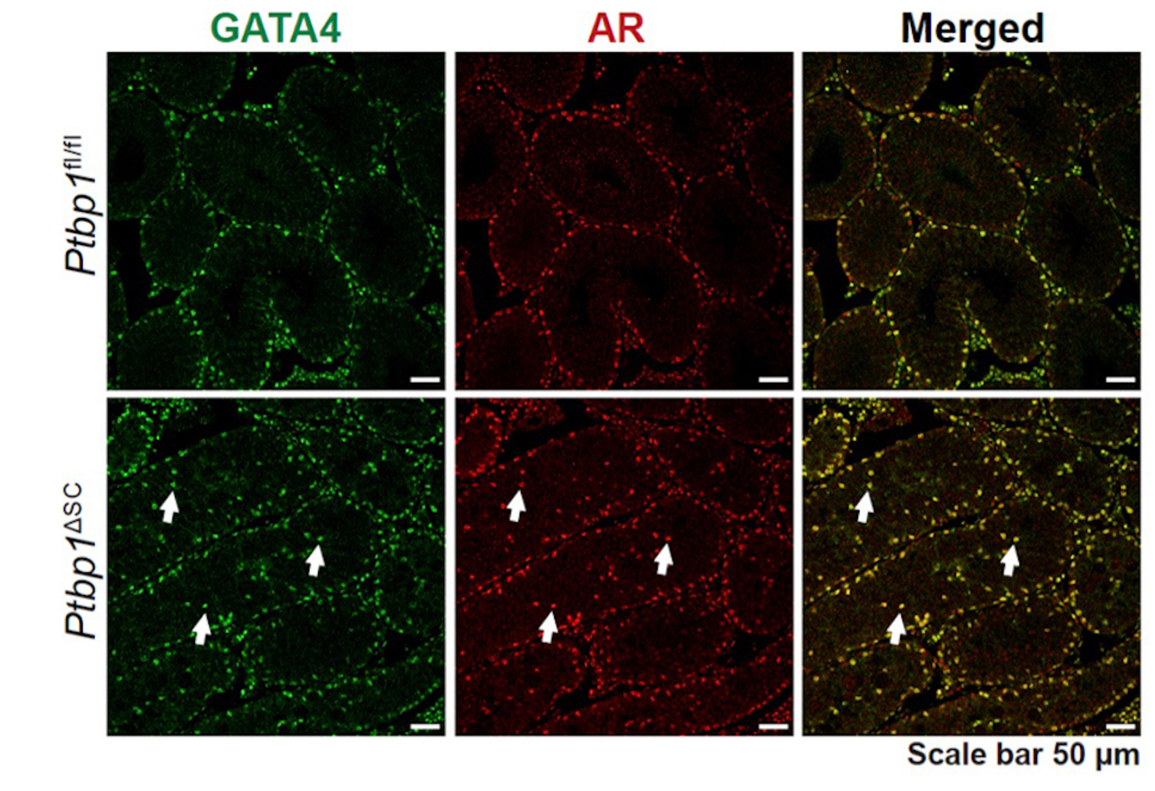
Maturation of Sertoli cells is not affected in *Ptbp1*^ΔSC^ mice. Double immunofluorescent staining using antibodies against GATA4 (to mark Sertoli cells) and androgen receptor (AR) (to mark mature Sertoli cells) shows the maturation of Sertoli cells in *Ptbp1*^fl/fl^ and *Ptbp1*^ΔSC^ mice. Arrows point to mislocalized Sertoli cells in *Ptbp1*^ΔSC^ mice.

**Supplementary Fig 5.**
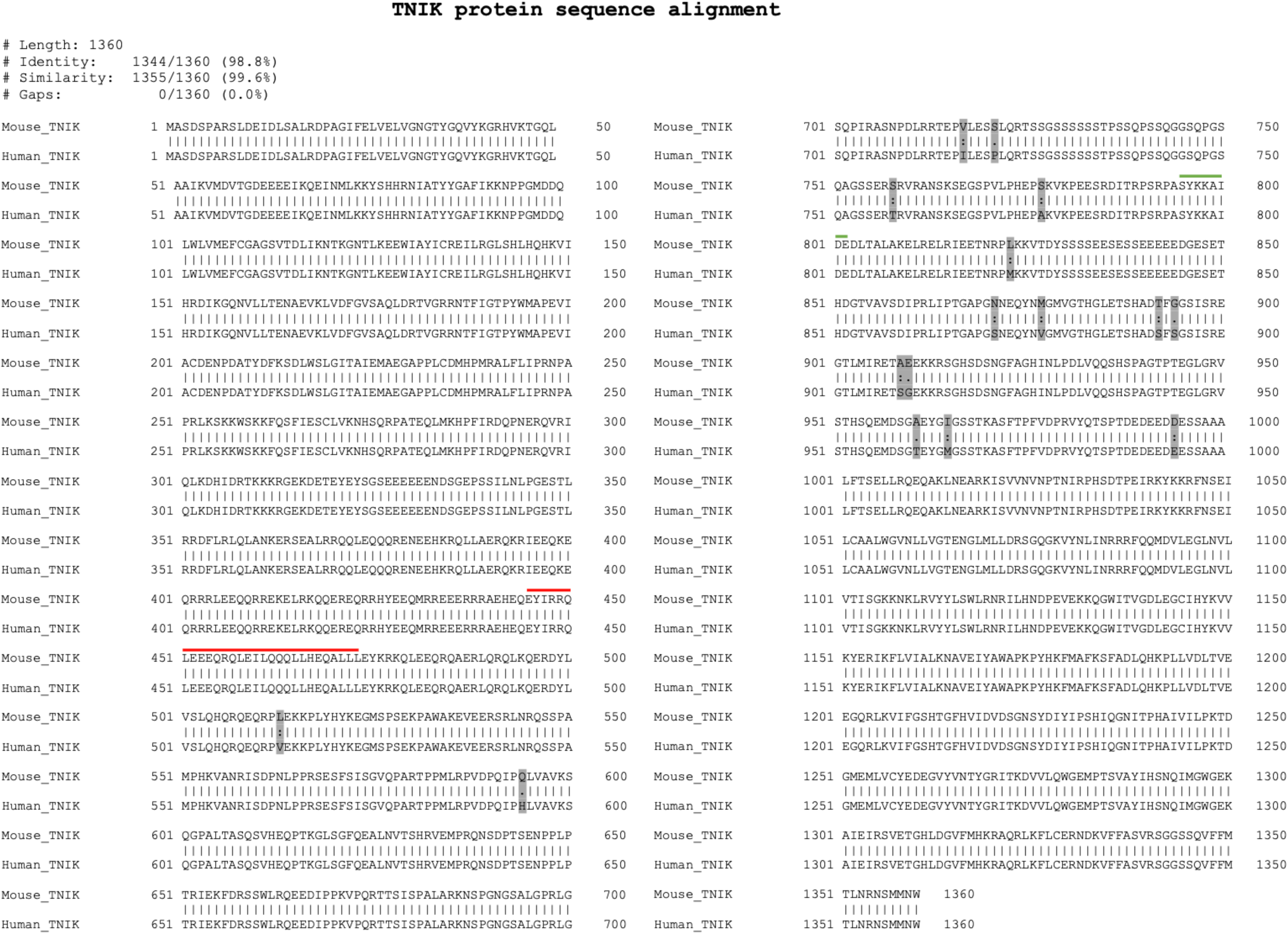
Alignment of Mouse and human Tnik protein sequences. Mouse and human TNIK proteins share a 99% similarity. Exon 14 encoded 29 amino acids are 100% identical (indicated by the red line above the sequence) between human and mouse TNIK. The green line above the sequence indicates exon 21 encoded amino acids.

**Supplementary Fig 6.**
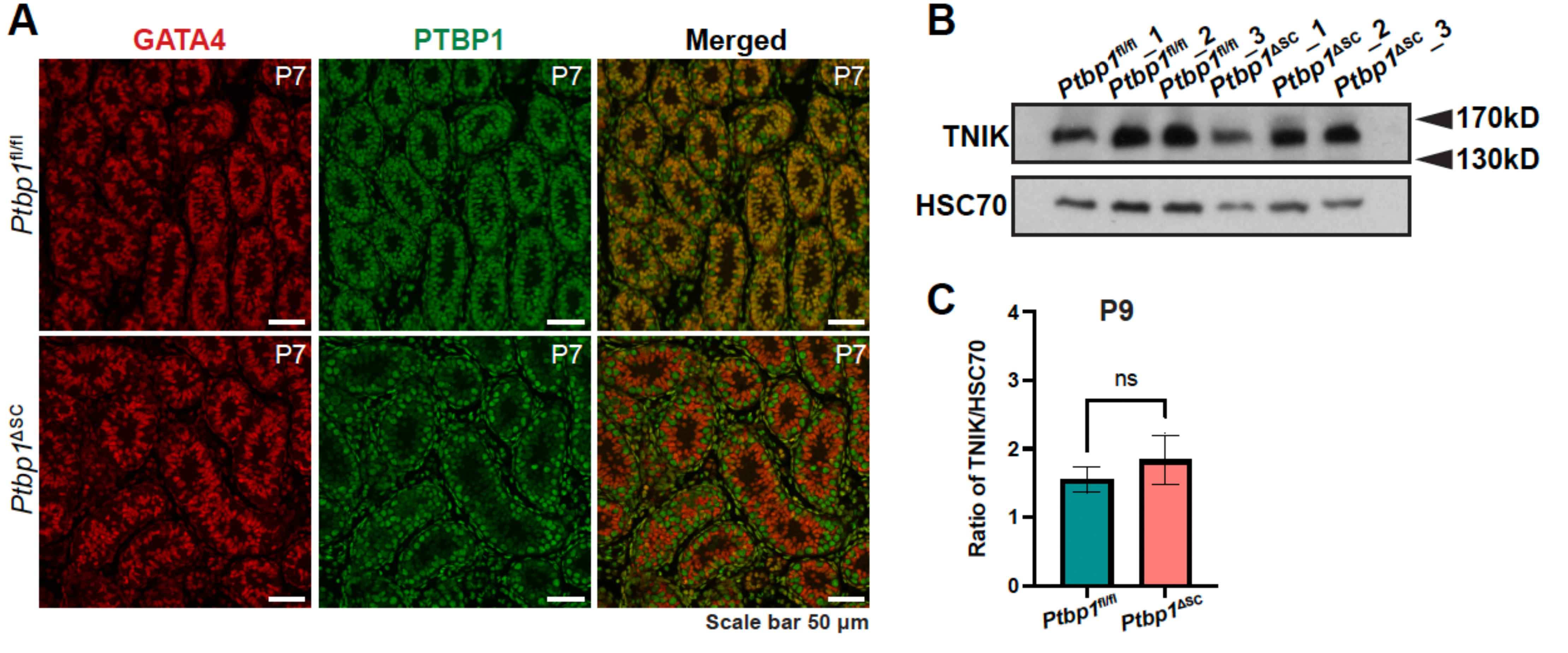
PTBP1 deficiency does not alter the protein level of TNIK. (**A**) Double immunofluorescence staining using antibodies against GATA4 and PTBP1 shows the efficient PTBP1 deletion in Sertoli cells at P7. (B-C) Western blotting result shows the total levels of TNIK protein in the testes of *Ptbp1*^fl/fl^ and *Ptbp1*^ΔSC^ mice at P9 are comparable. Quantification of the ratio of TNIK to HSC70 in the testis is shown in C. Mice used in the assay are littermates. Data are presented as mean± SD. ns, not significant.

**Supplementary Table 1.**
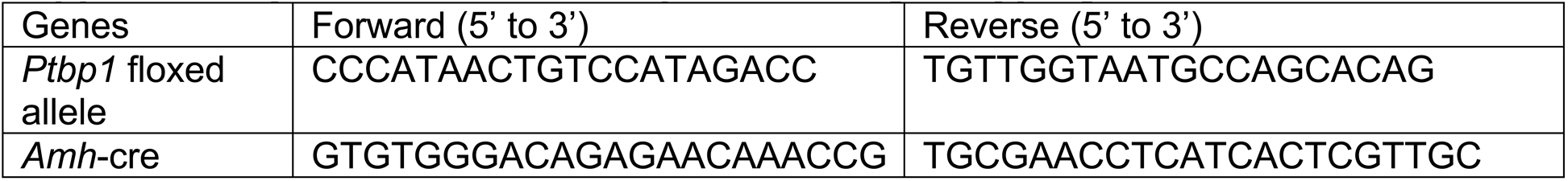
Primer sequences for genotyping.

**Supplementary Table 2.**
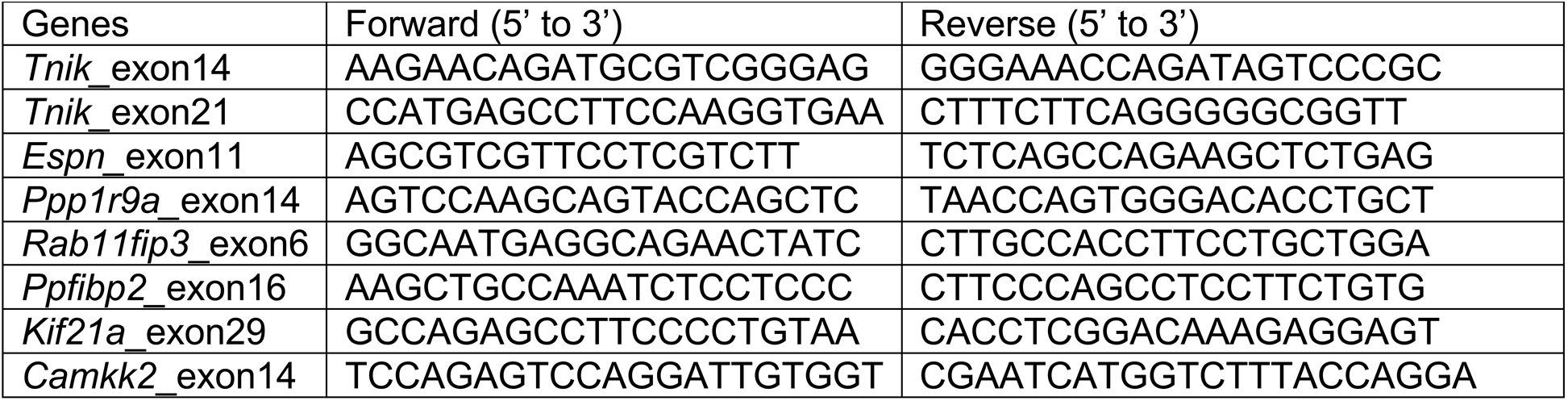
Primer sequences for splicing assay.

**Supplementary Table 3.**
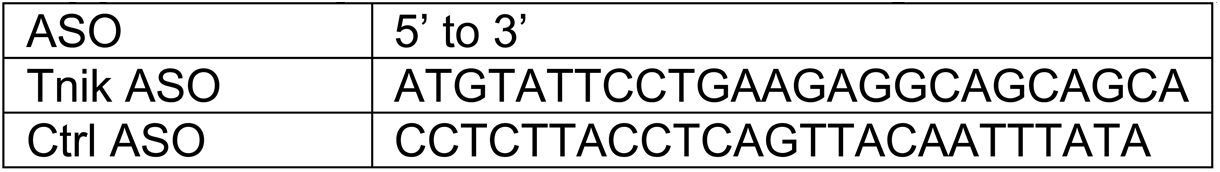
Antisense oligonucleotide (ASO) sequences.

